# Altered Proteasome Composition in Aging Brains, Genetic Proteasome Augmentation Mitigates Age-Related Cognitive Declines, and Acute Proteasome Agonist Treatment Rescues Age-Related Cognitive Deficits in Mice

**DOI:** 10.1101/2024.10.17.618893

**Authors:** Andrew M. Pickering

## Abstract

The aging brain experiences a significant decline in proteasome function, The proteasome is critical for many key neuronal functions including neuronal plasticity, and memory formation/retention. Treatment with proteasome inhibitors impairs these processes. Our study reveals a marked reduction in 20S and 26S proteasome activities in aged mice brains driven by reduced functionality of aged proteasome. This is matched by a decline in 20S proteasome but an increase in 26S proteasome. Our data suggests this may be a compensatory response to reduced functionality. By overexpressing the proteasome subunit PSMB5 in the neurons of mice, enhancing proteasome function, we slowed age-related declines in spatial learning and memory as well neuromuscular declines. We then showed acute treatment with a proteasome activator to rescue spatial learning and memory deficits in aged mice. These findings highlight the potential of proteasome augmentation as a therapeutic strategy to mitigate age-related cognitive declines.

## INTRODUCTION, RESULTS, DISCUSSION

Declines in proteasome function are a robust feature of aging reported in human T-lymphocytes (Ponnappan, Zhong et al. 1999), rodent heart, kidney, liver, lung, and muscle (Bardag-Gorce, Farout et al. 1999, Keller, Hanni et al. 2000). In the nervous system, proteasome activity decreases in the cortex, cerebellum, and spinal cord, but remains unchanged in the cerebellum and brain stem (Keller, Hanni et al. 2000). Similar declines are observed in the heads of fruit flies(Munkacsy, Chocron et al. 2019) and the brains of killifish (Kelmer Sacramento, Kirkpatrick et al. 2020). This decline correlates with increased levels of oxidized and polyubiquitinated proteins (Petropoulos, Conconi et al. 2000, Rai, Coleman et al. 2021). We examined proteasome activity and assembly in whole brains from young (12 month old mice), compared to old (22-26 month old mice) (***Fig. 1A***). Here we employed a fluorescent proteasome activity probe MV151 which selectively binds to active proteolytic subunits of proteasomes (Verdoes, Florea et al. 2006) and then separated proteasome forms by native-PAGE gel. We observed a significant decline in activities of both 20S and 26S proteasome forms with age (***Fig. 1A***). Notably although activities of both proteasome forms declined with age we observed a more pronounced 70% decline in 20S proteasome activity compared to a 50% decline in 26S proteasome activity resulting in a shift in proteasome activities with age towards 26S activity (***Fig. 1B***).

**Figure 1.**
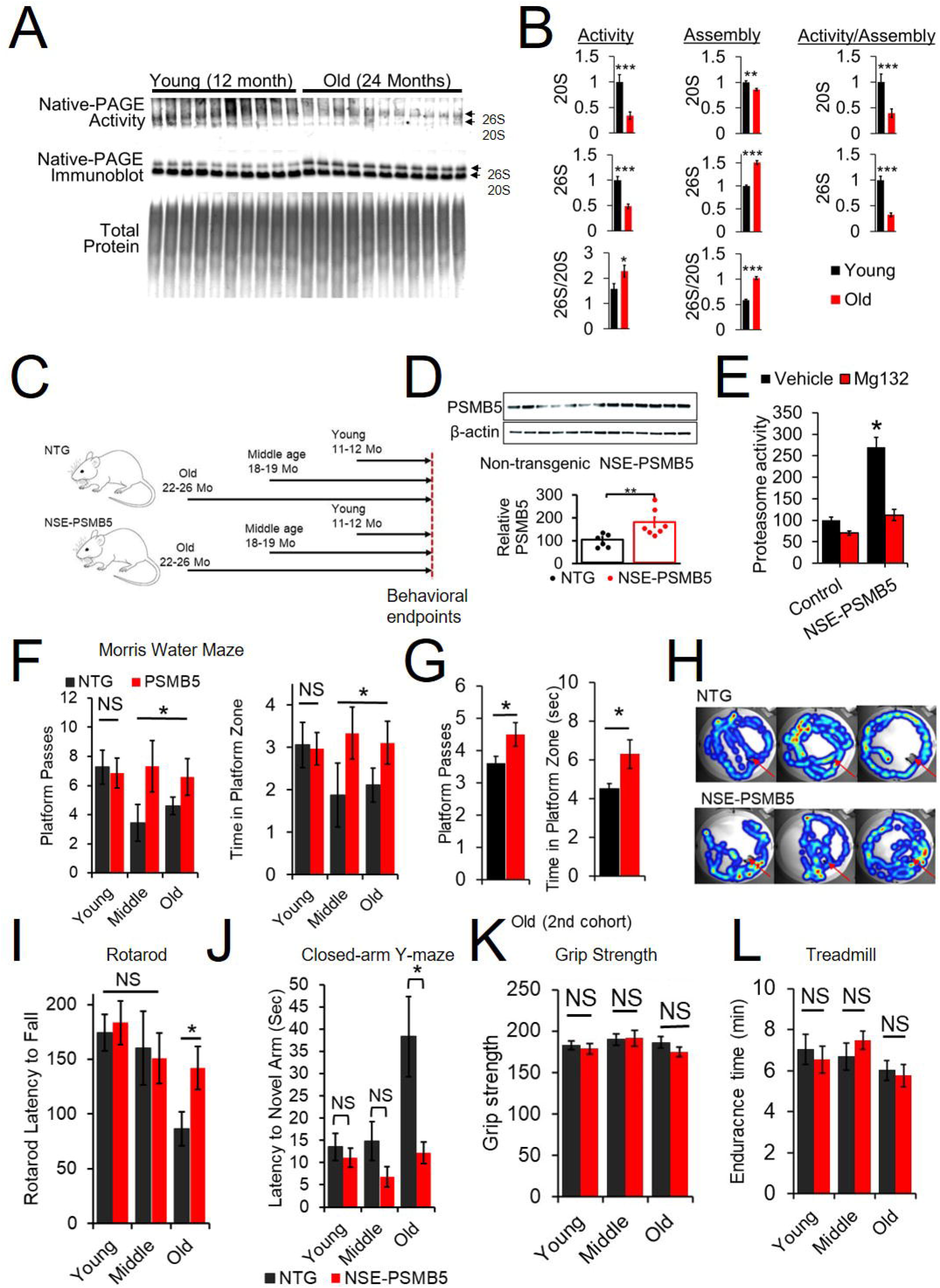
Neuronal PSMB5 overexpression delays age-related declines in cognitive tasks. (A) Native PAGE blot of whole brains from 10 young (12±1 Mo mice) and 11 Old (24±2 Mo mice), Top gel shows proteasome activity measured by Native PAGE gel of samples incubated with the MV151 fluorescent proteasome probe. This is a broad proteasome inhibitor binding to active centers and producing fluorescence. Middle gel shows a Native-PAGE immunoblot against the proteasome subunit PSMB5. Bottom gel shows an SDS-PAGE quantification of total protein by silver stain. (B) Quantification of Native PAGE activity and immunoblots. (C) Experiment design, 3 cohorts were staggered to reach young, middle and old age points then assayed. (D) Immunoblot of PSMB5 normalized to β-actin, brains dissected from 3month-old mice. NTG, N = 6-7; (E) Relative proteasome peptidase (chymotrypsin-like) activity, brains dissected from 3-month-old mice (N = 5 to 6). Replicate samples were incubated with the proteasome inhibitor MG132. (F) NSE-PSMB5 reduces age-related learning and memory deficits in Morris water maze. (G) Morris water maze in a second cohort of aged animals (H) Representative heatmaps of water maze probe trial, arrow shows where platform was prior to removal. (I) NSE-PSMB5 reduces age-related latency to fall in rotarod assay. (J) NSE-PSMB5 reduces latency to explore the novel arm in a closed arm Y-maze assay. (K) NSE-PSMB5 did not alter grip strength. (L) Treadmill maximum speed and endurance time declined with age, NSE-PSMB5 did not alter performance. Cohort 1 comprised 11 young NTG, 15 young NSE-PSMB5, 7 Middle age NTG, 3 Middle age NSE-PSMB5, 10 old age NTG, 9 old age PSMB5. Cohort 2 comprised 11 old NTG and 12 old NSE-PSMB5. *P < 0.05, NS = Not Significant.

We next examined proteasome assembly by Native-PAGE immunoblot. We saw a significant but small decline in 20S proteasome assembly with age. However strikingly we saw a pronounced and reproducible 50% increase in 26S proteasome assembly with age leading to an increase in 26S assembly compared to 20S assembly (***Fig 1B***). This finding was surprising as prior studies have reported a decline rather than an increase in 26S proteasome assembly with age (Vernace, Arnaud et al. 2007, Dasuri, Zhang et al. 2009). We hypothesize this discrepancy might be due to differences in the age of the ‘young’ animals. Prior studies employed 3 month old mice to as a representation of young animals(Dasuri, Zhang et al. 2009), while we employed 12 month old animals as our young animals. There is a substantial elevation in protein translation rates in mice in the first 3-6 months of life and we suggest there may be a commensurate increase in 26S proteasome in the first few months of life potentially explaining this discrepancy(Ward and Richardson 1991, Kim, Parker et al. 2023). When we examined the ratio of proteasome activity to abundance, we observed an about 70% decline in functionality of both 20S and 26S proteasome (***Fig. 1B***). This suggests that the decline in proteasome function with age is driven by a reduction in functionality of the proteasome rather than a decline in levels/assembly. We also suggest that the increase in 26S proteasome assembly we observe might represent a compensatory response to decreased proteasome functionality.

The proteasome plays a critical role in various neuronal functions including modulation of synaptic plasticity, dendritic spine growth, long term potentiation, memory formation and consolidation. In each of these cases treatment with proteasome inhibitors has produced deficits in these processes (Lopez-Salon, Alonso et al. 2001, Hamilton, Oh et al. 2012, Davidson and Pickering 2023). For this reason, we hypothesized that impairment in proteasome function in the aging brain may contribute to age-related cognitive declines. To test this hypothesis, we examined measures of cognitive performance in young, middle aged and old mice (12±1, 18±1 and 24±2 Mo respectively) in which we enhanced proteasome function through overexpression of the proteasome subunit PSMB5 (***Fig. 1C***).

Overexpression of the rate-limiting proteasome subunit PSMB5 boosts proteasome function and assembly in cell culture and invertebrate model organisms (Chondrogianni, Tzavelas et al. 2005, Chondrogianni, Georgila et al. 2015, Munkacsy, Chocron et al. 2019, Nguyen, Rana et al. 2019), extending lifespan in nematode worms (Chondrogianni, Georgila et al. 2015) and fruit flies(Munkacsy, Chocron et al. 2019, Nguyen, Rana et al. 2019). Overexpression of PSMB5 in the nervous system in flies delays age-related declines in olfactory aversion training assays (as a measure of learning and memory (Munkacsy, Chocron et al. 2019). We developed a mouse model with enhanced neuronal specific overexpression of PSMB5 (Chocron, Munkacsy et al. 2022) (***Fig. 1D***). This mouse displays increased proteasome activity in its brain (***Fig. 1E***), along with increased proteasome assembly(Chocron, Munkacsy et al. 2022). A more detailed characterization of this line is provided in our prior paper where we demonstrate protective capacity against AD pathology (Chocron, Munkacsy et al. 2022).

Evaluating spatial learning and memory via Morris water maze, we observed an age related decline in platform passes and time in the platform zone during our probe trial comparing young mice with middle and old aged mice. Combining our middle and old aged mice we observed a significant improvement both in platform passes and greater time in the platform zone in NSE-PSMB5 overexpression mice (***Fig. 1F***). Because of high variability in the water maze assay, to increase our confidence we repeated this assay in a second larger cohort of just old mice. This cohort, recapitulated our findings showing increased platform passes and time in the platform zone (***Fig 1G-H***). Additionally, we observed an age-related decline in neuromuscular function measured by rotarod which was slowed in NSE-PSMB5 overexpression mice (***Fig 1I***). Similarly, as another measure of spatial learning and memory, we observed reduced performance in a closed-arm Y-maze with age, which was improved in NSE-PSMB5 overexpression mice (***Fig. 1J***).

To test if the improvements in cognitive tasks we observed were produced by changes in physical function in our mice we evaluated grip strength and treadmill performance. We did not see any changes either with age or our transgene in grip strength (***Fig. 1K***). In contrast we observed an age-related decline in treadmill performance. Mice showed an age-related decline in maximal speed and endurance time on the treadmill. There was no significant impact from the transgene (***Fig. 1L***).

We next investigated if transient augmentation of proteasome function via treatment with a proteasome activating compound could rescue age-related cognitive deficits. We developed a set of proteasome activating small molecules which show a robust ability to enhance both 20S and 26S proteasome function *in vivo* and *in vitro*, a detailed characterization is reported in our prior publications (Gizynska, Witkowska et al. 2019, Osmulski, Karpowicz et al. 2020, Chocron, Munkacsy et al. 2022). We recapitulated our prior finding that our lead TAT1-DEN can enhance 20S proteasome function in the brains of mice under IP injection (***Fig. 2A-B***). Previous studies have shown acute treatment with proteasome inhibitors to produce deficits in long term memory formation and retrieval(Lopez-Salon, Alonso et al. 2001, Artinian, McGauran et al. 2008, Lee, Choi et al. 2008). Our running hypothesis is that age-related declines in proteasome function may produce similar deficits. We thus hypothesized that acute treatment with a proteasome activator might rescue age-related deficits in memory formation. To test this hypothesis, we employed young 12-month-old mice alongside old 24-month old mice each IP injected with our proteasome agonist.

**Figure. 2.**
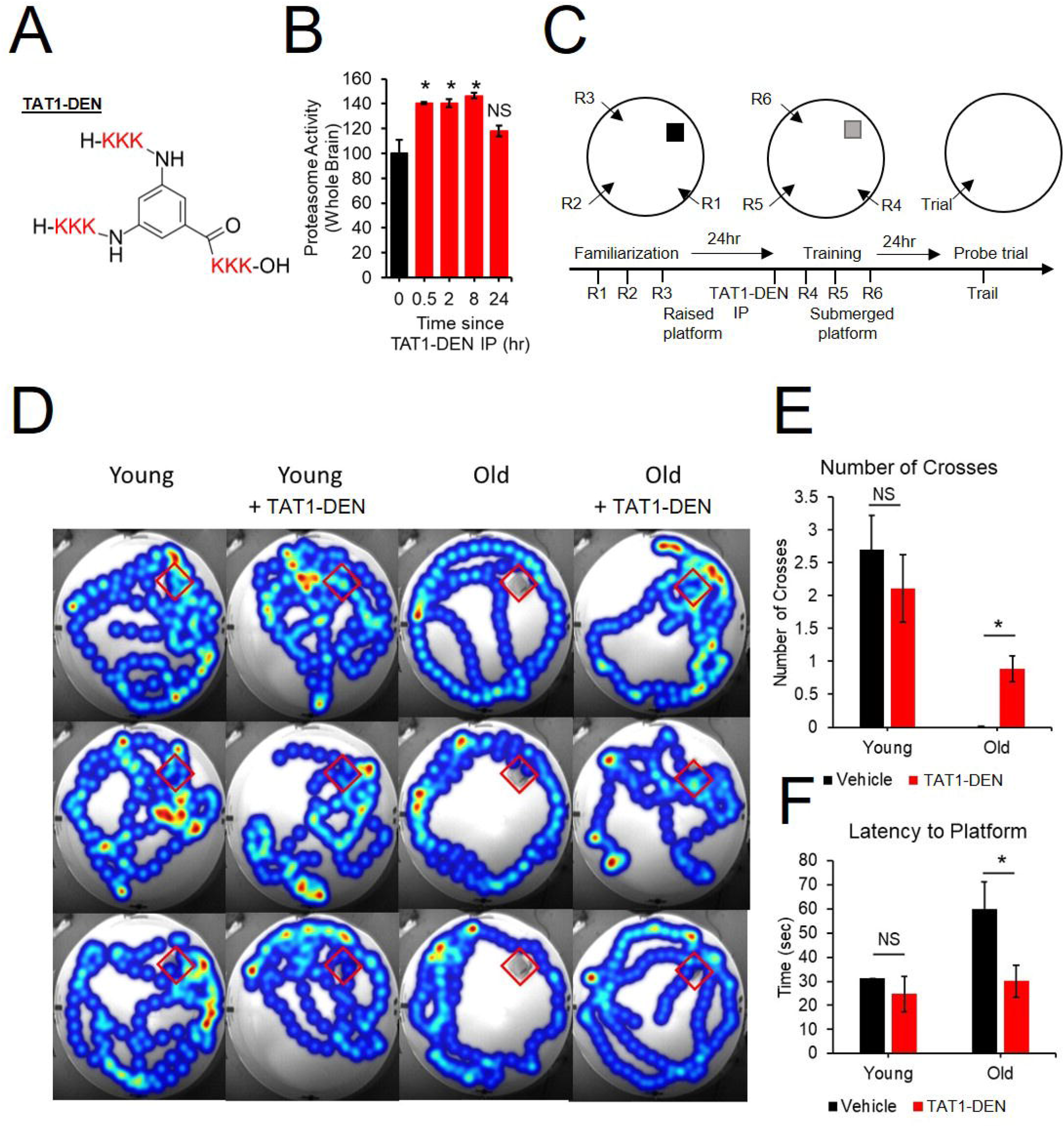
Acute treatment with proteasome agonist TAT1-DEN rescue age-related spatial learning and memory deficits. (A) Structural schematic of compound (B) Proteasome activity (suc-llvy-amc degradation) in whole brain lysate from animals IP injected with TAT1-DEN. Animals were sacrificed and lysates generated after 30min, 2hr, 8hr, and 24hr, N = 2-4. (C) Modified Water maze experiment design based on design in(Artinian et al., 2008). (D-F) Age reduces performance in Morris water maze, IP injection with TAT1-DEN rescues deficits. (D) representative heatmaps (3 per group) of probe trial, red boxes shows location of platform prior to removal. Background Photographs were taken at trial start prior to platform removal. recording was stopped once animals reached platform. (E) Number of Crosses of platform zone in probe trial. (F) Latency to platform zone in probe trial. N = 19 young control 18 Young TAT1-DEN treated, 14 old control, 13 old TAT1-DEN treated. *P < 0.05, NS = Not Significant.

Animals were evaluated for spatial learning and memory via a modified version the of Morris water maze. We employed a design similar to the design used by Artinian and colleagues(Artinian, McGauran et al. 2008) where they performed familiarization of animals to the maze on day 1, acquisition training on day 2 followed by injection of a proteasome inhibitor and a probe trial on day 3. In this design they reported treatment with a proteasome inhibitor to produce spatial memory deficits (Artinian, McGauran et al. 2008). In our design, animals were first acclimated to the maze using a raised platform at multiple entry locations on day 1. On day 2 animals received an IP injection of the proteasome activator TAT1-DEN then 3 training sessions at multiple entry sites with the platform submerged. On day 3 a probe trial was performed (***Fig. 2C***). We demonstrated that aged mice show deficits in memory formation showing reduced crosses of the platform zone and increased latency to the platform zone in the probe trial. Treatment with our proteasome activator produced no effect in young animals but significantly improved performance in aged animals (***Fig. 2D-F***).

In conclusion we show aging to be associated with significant declines in proteasome function driven by reduced functionality of the proteasome rather than changes in expression/assembly. We show that enhancing proteasome levels and assembly in the nervous system through overexpression of a rate limiting proteasome subunit can slow age-related decline in spatial learning and memory as well as neuro-muscular function. Physical function is unaffected. Finally, we show that acute treatment with a proteasome activator can rescue age-related deficits in spatial learning and memory.

## ACKNOWLEDGEMENTS

We thank Joddie Cropper who performed animal genotyping as well as husbandry and Danitra Parker who assisted with behavioural measures. Funding: This work was supported by the National Institute of Aging R56 AG061051 (to A.M.P.), National Institute of Aging R01 AG065301 (to A.M.P.), 2018 Glenn Foundation for Medical Research and AFAR Grants for Junior Faculty (to A.M.P.).

## DATA AVAILABILITY STATEMENT

The data that support the findings of this study are available from the corresponding author upon reasonable request.

## Notes

### Competing Interest Statement

A.M.P. is an inventors on a patent application related to this work filed by The University of Texas Health Science Center at San Antonio (HSC1567, filed 13 September 2019). The author declares that they have no other competing interests.

